# High-Throughput Quantification of miRNA-3′-Untranslated-Region Regulatory Effects

**DOI:** 10.1101/2024.12.05.626985

**Authors:** Stephen Mastriano, Shaveta Kanoria, William Rennie, Chaochun Liu, Dan Li, Jijun Cheng, Ye Ding, Jun Lu

## Abstract

MicroRNAs (miRNAs) regulate gene expression post-transcriptionally, primarily through binding sites in 3′ untranslated regions (3′ UTRs). While computational and biochemical approaches have been developed to predict miRNA binding sites on target messenger RNAs, reliable and high-throughput assessment of the regulatory effects of miRNAs on full-length 3′ UTRs can still be challenging. Utilizing a miniaturized and high-throughput reporter assay, we present a ‘pilot miRNA-targeting map’, containing 4,994 successfully measured miRNA:3′ UTR regulatory outputs by pairwise assays between 461 miRNAs and eleven 3′ UTRs. This collection represents a large experimental miRNA:3′ UTR dataset to date on a single platform. The methodology can be generally applied to studies of miRNA-mediated regulation of critical genes. We found that seedless sites can lead to substantial downregulation. We utilized this dataset in the development of a quantitative total score for modeling the total regulatory effects by both seed and seedless sites on a full-length 3′ UTR. To assess the predictive value of the total score, we analyzed data from mRNA expression and proteomics studies. We found that the score can discriminate the potent miRNA inhibition from the weak inhibition and is thus useful for quantitative prediction of miRNA regulation. The score has been added to the STarMir program of the Sfold package now available via GitHub at https://github.com/Ding-RNA-Lab/Sfold.

## Introduction

As a class of small non-coding RNAs, microRNAs (miRNAs) regulate diverse biological processes by post-transcriptionally modulating gene expression [1–3]. miRNAs bind to partially complementary sites on target messenger RNAs to downregulate target protein expression by destabilization of target mRNA and/or inhibition of protein translation. Experimental scientists working on miRNAs frequently face the question of which miRNAs regulate their genes of interest. This is often challenging because it is important to assess the level of miRNA-mediated regulation in the context of the whole gene, in addition to knowing that a miRNA can bind to a mRNA. miRNAs that can only weakly regulate targets are less interesting for further investigation.

Two approaches are often used to identify miRNAs that target genes of interest. Computational target predictions rely mainly on sequence information and known rules of miRNA:target recognition. It is widely perceived that computational algorithms have high levels of false-positive and false-negative rates [4,5] Additionally, these algorithms typically predict whether a miRNA can regulate a target gene, rather than the level of miRNA-mediated target regulation. The second approach is the biochemical purification of miRNAs and target binding sequences. For example, cross-linking and immunoprecipitation (CLIP)-based methodologies are powerful and capable of revealing miRNA-target binding events in cells [6–9]. However, it is impossible to deduce the functional output of these binding events. This is because CLIP methods do not inform the percentage of target messenger RNA occupied by a specific miRNA; it is also conceivable that binding by some miRNAs may not lead to effective functional repression of target protein production. As such, it is impossible for either approach to reliably assess the level of miRNA-mediated regulation. While one can estimate the level of miRNA-mediated regulation on predicted target genes via overexpressing or knocking out a miRNA followed by transcriptomic analyses, these transcriptomic level assays are prone to secondary and tertiary effects.

The high false rates of computational algorithms are likely due to an incomplete understanding of the miRNA-mediated target regulation. Some significant observations are summarized here. (1) Although miRNA binding sites have been experimentally identified in 3′ UTRs, coding sequences and beyond, it is believed that most functional regulatory output occurs through 3′ UTR binding sites [2]. (2) The 5′ region of the miRNA (also known as the “seed” region), particularly involving nucleotides 2-8, is critical for target recognition [2,10]. The Bartel laboratory has further classified seed-based interaction on target 3′ UTR into several categories. The presence of an “8mer” site is correlated with strong repression, “7mer” site with medium effect, whereas “6mer” site has weak to no effect [2,11]. (3) Other than seed-based interactions, seedless sites have also been observed to mediate miRNA binding on target 3′ UTRs [12–15], although the effect of such sites on target regulation is debated [8,14–16] and deemed very weak by some [17]. (4) The neighboring sequence content, the location of a miRNA binding site within 3′ UTR [6,18], and the structural accessibility of the target [19–22], can all influence miRNA binding and effectiveness.

In this study, we used a miniaturized cell-based 3′ UTR reporter assay to allow high-throughput quantification of miRNA-mediated regulation on 3′ UTR. It is important to note that our goal was to assay the effect from an abundant miRNA source in order to identify possible miRNA-mediated target regulation rather than physiological regulation. With this approach, we present an initial dataset (termed pilot miRNA-targeting map) that is comprised of 4,993 interactions for 11 3′ UTRs and 461 miRNAs, obtained on a single platform. We used this dataset to develop a quantitative score for modeling the total regulatory effects by both seed and seedless sites.

## Results

### Developing a high-throughput cell-based reporter assay to quantify miRNA:3′ UTR regulation

Luciferase reporter assay has long been a gold-standard to quantify miRNA-mediated regulation of 3′ UTR activity and to validate computational predicted miRNA:3′ UTR interactions. The basis of the assay is that miRNA-mediated regulation directly via motifs within a 3′ UTR should be recapitulated on a non-related gene in a heterologous cellular background. It has been demonstrated that miRNA-mediated repression is not strongly influenced by cellular background [16]. Consistent with this notion, for a vast majority of reported cases, luciferase reporter assays reveal the functional consequences of direct miRNA-target interaction. Compared to measuring messenger RNA or proteins in cells which are prone to secondary and tertiary effects, the 3′ UTR reporter assay will allow lower false-positive reporting, because only a limited number of protein-coding genes are known to regulate 3′ UTRs and that miRNAs do not often regulate each other.

We have previously shown that we can use 384-well-based luciferase assay to quantify miRNA-mediated regulation of TET2 3′ UTRs [23]. However, whether this approach is reliable and expandable to many 3′ UTRs beyond TET2 has not been demonstrated, which led us to further optimize and test this approach (Fig. 1). Specifically, we used 293T cells for the assay, which is a cell line that has been widely used for assaying miRNA:3′ UTR interactions (e.g., [24,25]). In addition, 293T cells support high levels of gene expression in transient transfection assays, allowing enough signals for detecting luciferase activities.

**Figure 1.**
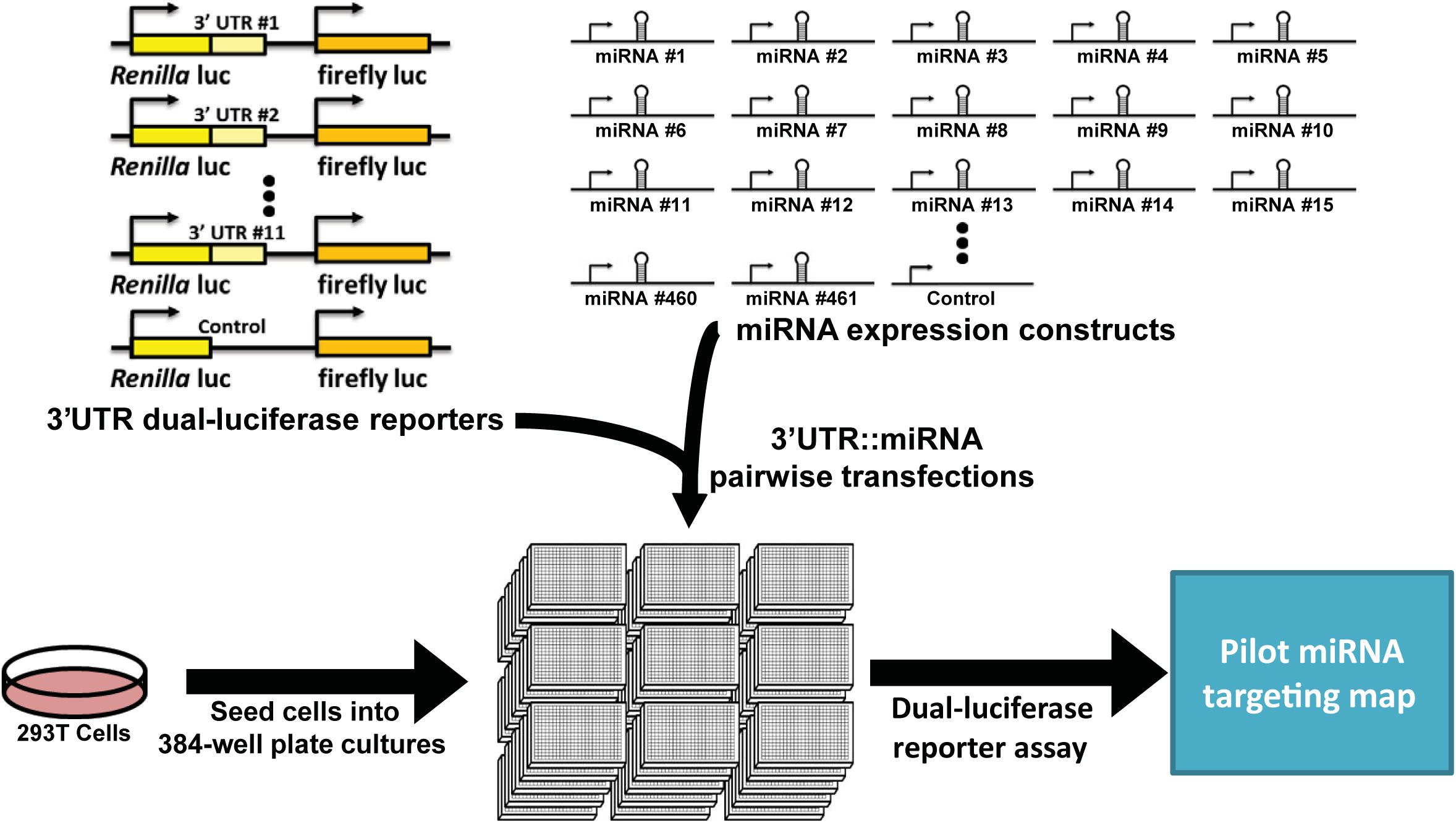
Schematic of a high-throughput 3′ UTR reporter assay and the generation of the pilot miRNA-targeting map. A schematic for the high-throughput 3′ UTR luciferase assay was used to generate the pilot miRNA Target Map, in which 461 miRNA expression constructs were assayed pairwise with eleven 3′ UTR dual-luciferase reporters in 293T cells.

Indeed, using this 384-well format, we were able to capture the know regulation of miR-25 suppressing KLF4 levels [26] using a KLF4 3′ UTR reporter. This is a dual luciferase reporter with the KLF4 3′ UTR regulating *Renilla* luciferase activity, whereas firefly luciferase activity serves as an internal control. We co-transfected the KLF4 3′ UTR reporter with a miR-25 expression vector, into 6-well plate, 96-well plate and 384-well plate wells. In each case, we observed ∼2-fold suppression of 3′ UTR activity (Fig. 2a), confirming literature findings. Importantly, the level of suppression observed in 384-well plates was similar to that observed in 6-well plates (Fig. 2a), indicating that the 384-well format can accurately capture miRNA-mediated 3′ UTR regulation.

**Figure 2.**
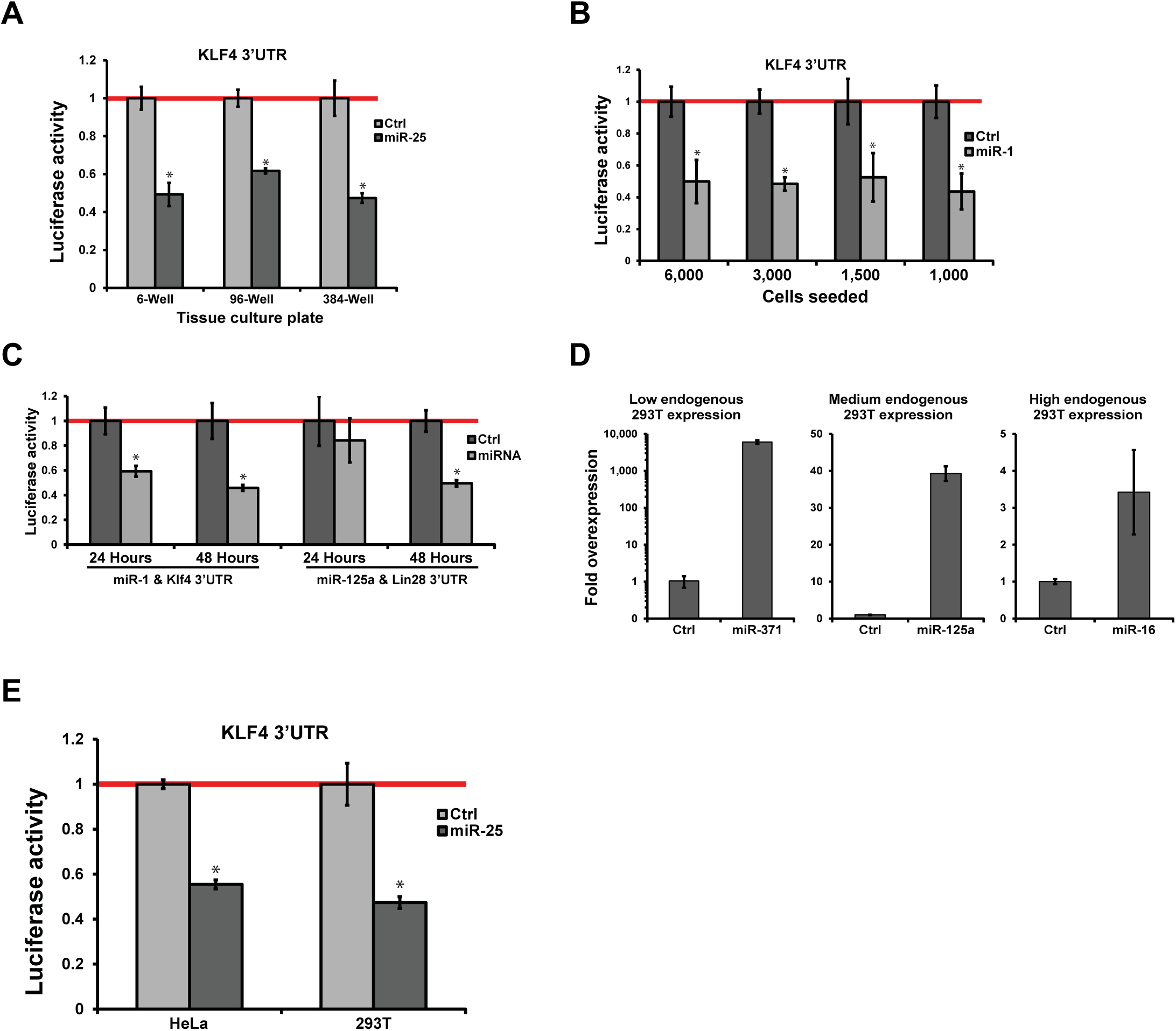
Assay parameter optimization for the high-throughput 3′ UTR reporter assay. **(a)** Down regulation of the Klf4 3′ UTR luciferase reporter by miR-25 in 6-, 96-, and 384-well plates was determined. 293T cells were transfected with a control miRNA construct (Ctrl) or miR-25. Normalized luciferase activities are plotted with one (red line) indicating no change. Error bars represent standard deviations. N = 4. **(b)** The seeding density of 293T cells was evaluated using Klf4 3′ UTR reporter assayed with miR-1. The numbers of 293T cells seeded per well in 384-well plates are indicated. Error bars represent standard deviation. N = 4. **(c)** Luciferase activity reading at 24 and 48 hours post-transfection was evaluated for miR-1-mediated suppression of KLF4 3′ UTR and miR-125a-mediated suppression of LIN28 3′ UTR. Normalized luciferase activities are shown. Error bars represent standard deviation. N = 4. **(d)** MiRNA constructs for miR-371, miR-125a, and miR-16, representing low, medium and high endogenous levels, were evaluated for overexpression in 293T cells 48 hours post transfection. A control miRNA construct (Ctrl) was used for comparison. MiRNA expression was determined by qRT-PCR. Representative data are shown. Error bars represent standard deviation. N = 3. **(e)** The regulation of miR-25 on KLF4 3′ UTR was evaluated in 293T and HeLa cells in comparison with a control miRNA construct. Error bars represent standard deviation. For HeLa, N=3; For 293T, N=4. For all sub-figures, *p<0.01.

Next, we optimized two key parameters of the assay, namely the number of cells to plate in each 384-well plate well, and the time to read out luciferase activity after transfection. Varying cell densities were tested using the known targeting of miR-1 on KLF4 3′ UTR [18]. All densities generated similar levels of down regulation for the KLF4 3′ UTR reporter by miR-1 (Fig. 2b). However, plating 3,000 cells per well was deemed optimal because lower cell densities produced weaker luciferase activity readings and thus higher variation (Fig. 2b), whereas higher density leads to overcrowding of cells in wells. We then tested two time points for reading out luciferase activity at 24 or 48 hours post transfection. Again, using the known targeting of miR-1 on KLF4 or miR-125a on LIN28, we determined that the 48-hour time point produced better results (Fig. 2c). Of note, cells became over-confluent at day 3 post-transfection. These experiments established the optimal condition for our 384-well reporter assays.

We had three considerations regarding the use of 293T cells as an assaying cell type. First, can we achieve exogenous overexpression for miRNAs endogenously expressed in 293T cells? Based on miRNA expression in 293T cells, we selected miR-371, miR-125a, and miR-16, with low, medium and high endogenous expression, respectively. Quantitative RT-PCR analyses indicate that all three miRNAs can be overexpressed, with the exact fold overexpression dependent on the endogenous basal levels of the corresponding miRNAs (Fig. 2d). Of note, the levels of overexpression are underestimated, because transfection efficiency is below 100%. Second, can we observe repression by miRNAs with high endogenous expression? Our profiling data indicate that miR-25 is among the most abundant endogenous miRNAs in 293T cells. Nevertheless, we were able to observe repression by this miRNA on KLF4 3′ UTR (Fig. 2a). Third, can 3′ UTR regulatory relationships revealed in 293T cells be observed in other cell types? For this purpose, we tested the repression of miR-25 on the KLF4 3′ UTR in HeLa cells. Unlike 293T cells, 384-well format for HeLa cells yielded luciferase signals too weak to be quantifiable. We thus compared data from HeLa cells assayed in 96-well plates, which showed similar levels of repression as observed in 293T cells (Fig. 2e) and is consistent with existing data supporting that different cellular backgrounds do not significantly impact miRNA-mediated gene regulation [16]. We will further examine this point in later sections.

Taken together, we established experimental conditions to reliably quantify miRNA-mediated repression of 3′ UTRs using 384-well-based high-throughput reporter assays.

### Generation of the pilot miRNA-targeting map

With a suitable tool in hand, we set out to quantify a large number of miRNA:3′ UTR pairs. We cloned eleven 3′ UTR reporters, representing both human (EIF2C2, DICER1, POU5F1, SOX2, KLF4, MYC, NANOG, LIN28, TP53 and MYB) and mouse (Bmi1) genes. This list of genes includes transcription factors, RNA binding proteins, epigenetic regulators, and cancer-related genes. For miRNAs, we used an expression library that we generated [23,27], containing 461 individual miRNA overexpression constructs. Each 3′ UTR reporter was assayed pairwise with each miRNA construct in quadruplicates (Fig. 1), generating a total of 31,249 data points (including replicates and controls), after filtering out wells with poor luciferase readings. We term this dataset the ‘pilot miRNA-targeting map’.

To ensure comparability of data across different 384-well plates, we built two types of in-plate controls, also in quadruplicates. These controls included control 3′ UTR vector with each of the miRNAs assayed on the plate (including miRNA vector control), and each of the 3′ UTRs assayed on the plate (including 3′ UTR vector control) with a control miRNA vector. We utilized the two types of controls when normalizing the luciferase activity, represented by the *Renilla* versus firefly ratio, so that a normalized value of 1 represents no regulation, and numbers below 1 representing downregulation (see Methods for details). We noticed that it is important to normalize with both types of controls, as the full dataset becomes much tighter after normalization, resembling a Gaussian distribution (Fig. 3a), which is expected because most miRNA:3′ UTR pairs will not have regulatory effects. This concept is further supported by the fact that with each additional control, there was a stepwise shrinking of overall data variation in the dataset.

**Figure 3.**
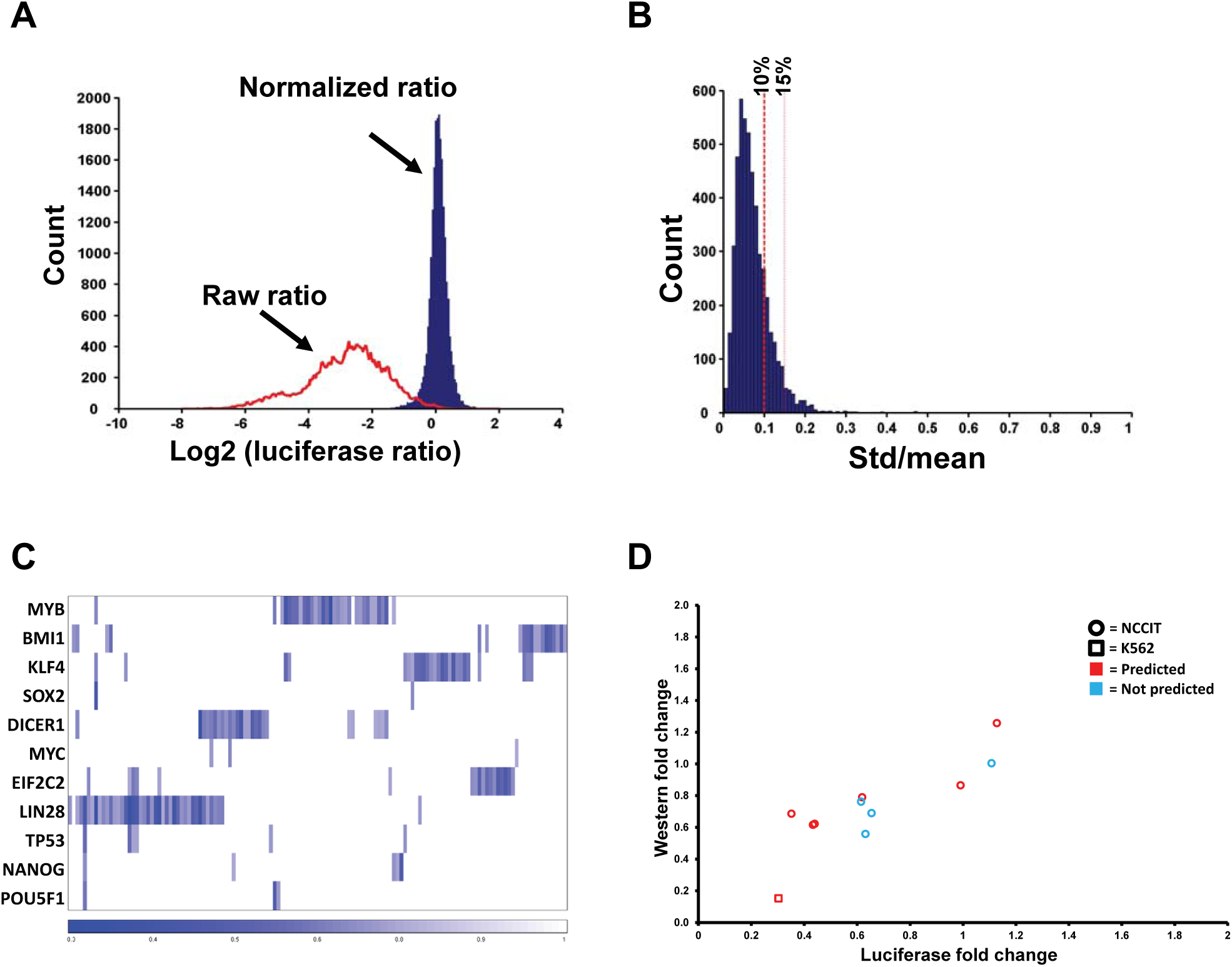
The pilot miRNA-targeting map. **(a)** Data normalization strongly reduced data variation. Raw and normalized *Renilla* vs firefly luciferase activities were plotted in a histogram. Note the shrinking of variation and near normal distribution of the normalized data. **(b)** Most assays in the pilot miRNA-targeting map have low levels of assay variation. A histogram is shown for standard deviation versus mean for the 4,994 miRNA:3′ UTR pairs assayed. Red dashed lines indicate 0.1 and 0.15 of standard deviation versus mean values. **(c)** A heatmap is shown to reflect the mean values of the 181 miRNA:3′ UTR pairs that resulted in ≥25% downregulation, with rows representing 3′ UTRs and columns representing miRNAs. Data were filtered to retain only those miRNAs that resulted in ≥25% downregulation for any of the assayed 3′ UTRs and were organized by hierarchical clustering afterward. Deeper blue reflects stronger suppression. MiRNA:3′ UTR pairs that did not result in ≥25% downregulation is set to 1 and represented by white color. **(d)** Eleven miRNA:3′ UTR pairs were evaluated on the level of endogenous protein. Corresponding miRNAs were transduced into NCCIT cells or K562 cells, and the corresponding endogenous protein changes were determined by western blot analysis. The mean luciferase fold changes from the pilot miRNA-targeting map were plotted against protein level fold change. Western data reflect averages of two independent biological replicates. Circles indicate analysis in NCCIT cells; squares indicate K562 cells. The color red indicates interactions predicted by TargetScan, PITA and mirSVR; blue indicates those interactions that were not predicted.

To evaluate the quality of our data, we next examined the variation of each assay. Among the 4,994 successfully measured miRNA:3′ UTR pairs, 79.2% showed a standard deviation ≤ 10% of the mean, and another 15.5% with a standard deviation between 10% and 15% of the mean (Fig. 3b). Taken together, the data above showed that we successfully generated the pilot miRNA-targeting map of overall good technical quality.

Among the 4,994 miRNA-3′ UTR pairs in the pilot miRNA targeting map, we identified 181 pairs with ≥25% down regulation (Fig. 3c). We set 25% downregulation as a main cutoff for two reasons. First, those repressive relationships of less than 25% will be less likely of interest to experimental scientists. Second, small levels of downregulation are more prone to experimental noise. Among the 181 downregulation pairs, at least 20 pairs (11%) were demonstrated previously, indicating that our assay captured known miRNA-mediated target regulation.

We compared our data to a biochemical binding dataset obtained from 293T cells using the CLASH method [8]. We chose CLASH data because it utilizes ligation to create miRNA-target chimeras which avoids the ambiguity of assigning miRNAs to target regions with other CLIP datasets. Due to the limited coverage of CLASH reads and possibly low expression of some associated genes in 293T cells, only seven miRNA:3′ UTR pairs overlapped between the CLASH data and our pilot miRNA-targeting map. Other than two pairs with a mild suppression, the other five pairs did not show downregulation in our data, supporting that not all CLASH sites result in functional suppression on 3′ UTRs. A much larger overlapping dataset would be needed to accurately estimate the percentage of functional CLASH sites.

Since we used 293T cells to generate the miRNA targeting map, we asked whether our data can reflect miRNA-target regulation in specific biological contexts other than 293T cells. To address this question, we examined eleven miRNA-3′ UTR pairs, including predicted pairs and those not predicted by any of three algorithms: TargetScan [11], PITA [21] and mirSVR [28] (Fig. 3d). These eleven pairs include ten miRNA targets expressed in human embryonic carcinoma cell line NCCIT. In addition, one pair involving MYB was examined in human hematopoietic cell line K562. We transduced corresponding miRNAs into relevant cell lines and measured endogenous protein expression by western blot.

Specifically, the endogenous protein levels of LIN28, POU5F1, SOX2, and NANOG were assayed in NCCIT, and MYB in K562 cells. Careful quantification of endogenous protein levels after miRNA expression revealed a significant correlation between western data and luciferase reporter assay data (Fig. 3d; R=0.845, p value=0.001). Noticeably, our data also indicate that multiple unpredicted miRNA:3′ UTR relationships can be validated on endogenous protein level (Fig. 3d). Due to the lack of seed sites in these miRNA:3′ UTR pairs and the difficulty in selection among many putative seedless sites in these 3′ UTRs for mutational analysis, we cannot exclude the possibility that such non-predicted regulation are through indirect effects. Nevertheless, our luciferase data indicate that our luciferase data can reflect *bona fide* miRNA:3′ UTR regulation, either directly or indirectly. The data above demonstrate that the miRNA-targeting map can reflect miRNA:3′ UTR regulation even though the target is expressed in a tissue specific context. Our data also suggest that the false-positive rate of the miRNA-targeting map is low, given that many of the regulatory relationships can be observed on the levels of endogenous proteins.

Among all miRNA-3′ UTR pairs for which targets were downregulated, 60.1% of the pairs only have seedless sites in the 3′ UTR; for ≥10% downregulation, 47.1% of the pairs only have seedless sites in the 3′ UTR; for ≥20% downregulation, 36.8% of the pairs only have seedless sites in the 3′ UTR; for ≥25% downregulation, 27.4% of the pairs only have seedless sites in the 3′ UTR. These suggest that seedless sites can lead to substantial target downregulation in the miRNA-targeting map.

### Development of quantitative score

We reasoned that we could use these data to develop computational scores for the quantitative prediction of miRNA-mediated regulation of full length 3′ UTRs. We used a linear weighted score for which the weighting coefficients were assigned to various site types (See Methods). To estimate the site type coefficients α_site-type_, we first used all available seed and seedless sites and performed the calculation as described in the Methods. However, the estimates did not meet the constraint a_8mer_ > a_7mer_ > a_6mer_ > a_seedless_. We reasoned that the data might contain too much noise, particularly due to the inclusion of weak miRNA binding sites. The logistic probability [22] can be used to filter out weaker sites with a low logistic probability. Specifically, we applied logistic probability thresholds to both seed sites and seedless sites, filtering the data to obtain a representative set of combinations based on the corresponding thresholds. We considered thresholds ranging from 0.0 to 0.90, with an increment of 0.05, for the logistic probability values. Among a total of 324 threshold pairs, we found two pairs that yielded efficacy parameter estimates satisfying the above constraint. The first pair is 0.75 for seed sites and 0.55 for seedless sites. The second pair is 0.75 for seed sites and 0.6 for seedless sites. Since the two sets of estimates produced highly similar results and the 0.55 threshold for the seedless sites includes more data, we preferred the estimates for the first pair of thresholds. Given those thresholds, the estimates of a_8mer_, a_7mer_, a_6mer_ and a_seedless_ are 0.0198, 0.0133, 0.00658 and 0.00428, respectively, with the intermediate statistics in the calculation presented in Table 1.

**Table 1:** Calculation of site-type specific efficacy.

| No. of 8mer sites | No. of 7mer sites | No. of 6mer sites | No. of seedless sites | RvF mean | No. of 3'UTR:miRNA pairs | Efficacy |
| --- | --- | --- | --- | --- | --- | --- |
| 28 | 0 | 10 | 699 | 20.39 | 24 | 0.01980 |
| 0 | 19 | 6 | 591 | 14.18 | 17 | 0.01327 |
| 0 | 0 | 76 | 1389 | 49.55 | 56 | 0.00658 |
| 0 | 0 | 0 | 7279 | 342.83 | 374 | 0.00428 |

With the estimates of the site type coefficients, we can compute the quantitative score for any miRNA:target pair as proposed in the Method. To assess the predictive value of the score, we analyzed high throughput mRNA and protein expression datasets from miR-1, miR-124 and miR-181a overexpression studies [29]. For all the down-regulated (mRNA/protein fold change less than or equal to 0) targets in the overexpression cell line, we compiled 8mers, 7mers, 6mers and seedless miRNA binding sites. For each miRNA target pair, in addition to score_total_, the score for seed and seedless sites combined, we also computed score_seed_, the component of the total score for seed sites, and score_seedless_, the component of the total score for seedless sites (score_total_= score_seed_ + score_seedless_). A component score facilitates evaluation of contribution of seed or seedless sites. For example, scores for NM_000018 are shown in Table 2.

**Table 2:** An example of score calculation for NM_000018.

| Target | miRNA | No. of 8mer sites | No. of 7mer sites | No. of 6mer sites | No. of seedless sites | Score<sub>seed</sub> | Score<sub>seedless</sub> | Score<sub>total</sub> |
| --- | --- | --- | --- | --- | --- | --- | --- | --- |
| NM_000018 | hsa-miR-1 | 0 | 0 | 2 | 22 | 0.00444 | 0.04372 | 0.04816 |
| NM_000018 | hsa-miR-124 | 0 | 1 | 1 | 131 | 0.00587 | 0.26711 | 0.27298 |

We computed the total and the two component scores for 2,406 targets down-regulated at the mRNA levels and 1,635 targets down-regulated at the protein level. To compare the scores between the subset with the highest and lowest level of downregulation, we applied various thresholds for the log fold change of the mRNA or the protein expression levels. For the set of targets with highest downregulation, the log fold change thresholds were −1.0, −0.5, −0.4, −0.3 (i.e., the set is defined by values below the threshold). For the lowest downregulated set of targets, the thresholds wereand −0.05, −0.10, and −0.20 (i.e., values between the threshold and 0).

We first analyzed mRNA expression data. For threshold pairs of −1.0 and −0.05, the average score_seed_ for the set of targets with mRNA fold change values < −1.0 is 0.0205, which is significantly higher than the average of score_seed_ of 0.0087 for the set of targets with mRNA fold change values between −0.05 and 0 (*p*-value of 1.3 x 10^−7^). For the second threshold pairs of −0.5 and −0.05, the averages of score_seed_ are 0.0131 and 0.0087, with the difference being significant (*p*-value of 2.1 x 10^−5^). For the third threshold pairs of −0.4 and −0.10, the averages of score_seed_ are 0.0117 and 0.008, with a p-value of 4.9 x 10^−6^ for the difference. For the last threshold pairs of −0.3 and −0.20, the averages of score_seed_ of 0.0107, is significantly higher than 0.0082 (p-value of 9.9 x 10^−6^). These data are summarized in Fig. 4a.

**Figure 4.**
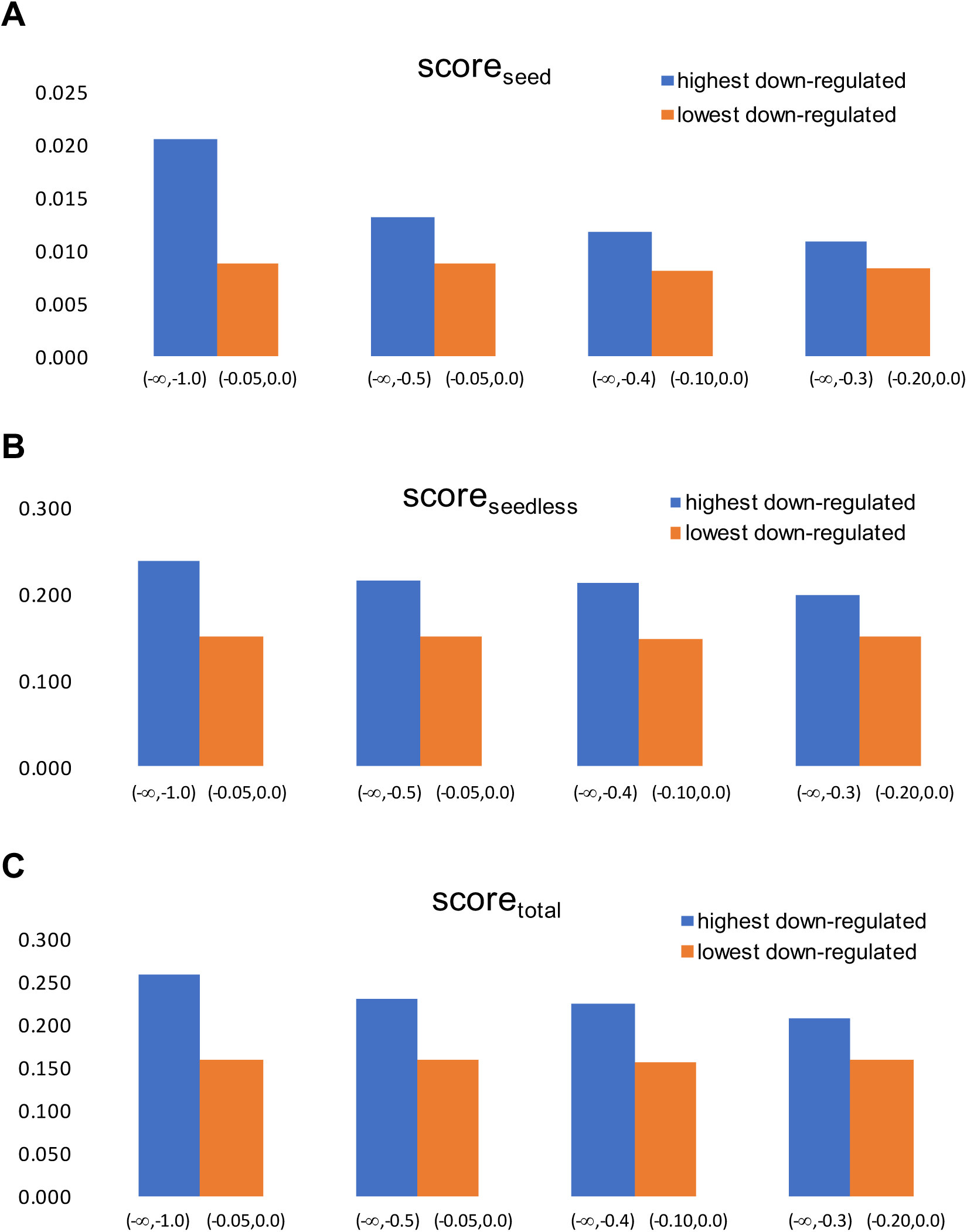
Validation of score by mRNA expression data. For each of the seed component score **(a)**, the seedless component score **(b)** and the total score **(c)**, the genes with the highest down-regulation in mRNA levels have significantly higher scores than those with lowest levels of down-regulation.

Similarly, for each of the threshold pairs above, the average score_seedless_ for the set of highest downregulation is significantly higher than that for the corresponding set of targets with lowest downregulation (Fig. 4b). E.g., for the threshold pair of −0.5 and −0.05, the difference between 0.2147 and 0.1497 has a *p*-value of 7.32 x 10^−9^. The same statistical trends were also observed for score_total_ (Fig. 4c). E.g., for threshold pair of −0.5 and −0.05, the difference between 0.2278 and 0.1584 has a *p*-value of 3.4 x 10^−9^. For the protein expression data, we also found that the average for each of the three scores is significantly higher for the set of targets with the highest levels of downregulation (*p*-value < 1.0 x 10^−15^ for all three scores; Fig. 5).

**Figure 5.**
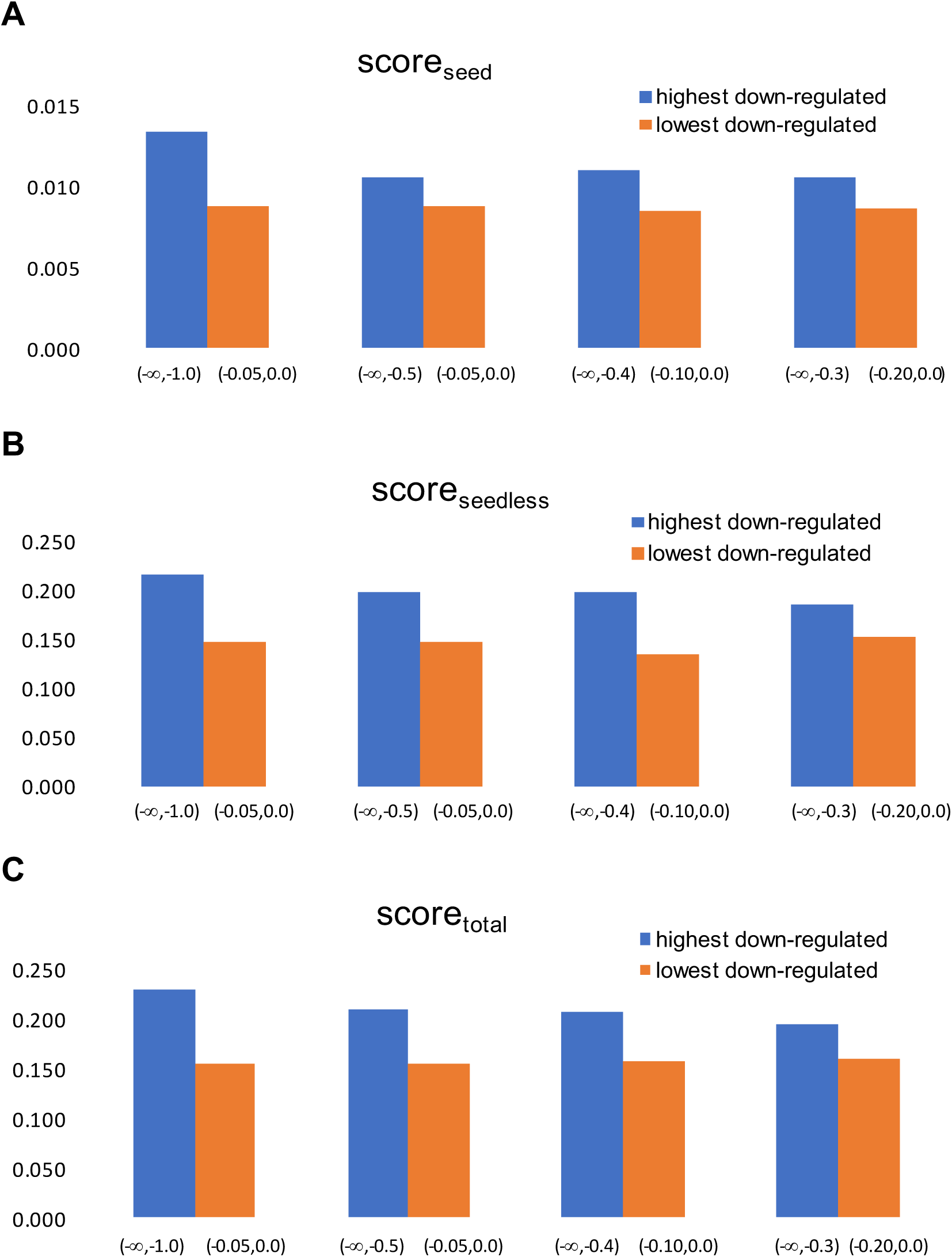
Validation of score by protein expression data. For each of the seed component score **(a)**, the seedless component score **(b)** and the total score **(c)**, the genes with highest down-regulation in protein levels has a significantly higher score than those with lowest levels of down-regulation.

The scores have been implemented in the STarMir program [30] of the Sfold package. In addition to web service (http://sfold.wadsworth.org/cgi-bin/starmirtest2.pl), STarMir is now freely available through Github at https://github.com/Ding-RNA-Lab/Sfold.

## Discussion

In this study, we present a pilot miRNA-targeting map based on a 384-well-plate high-throughput 3′ UTR reporter assay to quantify miRNA mediated regulation. This method can be particularly useful for experimental scientists to quickly identify miRNAs that regulate genes of interest, and to focus on those with stronger regulatory relationships. As an example, we have used this method to identify previously unappreciated oncogenic miRNAs by focusing on miRNAs that strongly regulate TET2, an epigenetic enzyme important in hematopoiesis [23]. It is important to note that, according to our data, miRNA regulation of a 3′ UTR does not necessarily imply physiological regulation which is dependent on the endogenous miRNA expression levels in specific cell types.

The pilot miRNA-targeting map contains quantitative regulatory outcomes from 461 miRNAs interacting with 11 3′ UTRs. This dataset differs from measuring genome-wide RNA or protein perturbations when a single miRNA is overexpressed or knocked out. It is conceivable that significant levels of secondary and tertiary effects on gene expression, even at early time points after miRNA perturbation, will increase the noise in the effort to decipher miRNA-targeting relationships. In contrast, by directly measuring 3′ UTR activity, we can avoid much of the secondary and tertiary effects, because there are relatively limited mechanisms that can influence 3′ UTR activity *in vitro*. Furthermore, when miRNAs are overexpressed, we rarely observe significant changes in endogenous miRNA expression. Our data is also different from measuring miRNA-target binding using biochemical approaches such as CLIP-based methods [6–9]. Such biochemical methods, although highly useful to inform in vivo miRNA binding on targets, do not necessarily indicate functional miRNA-based regulation. We observed that not all CLASH-identified miRNA:3′ UTR pairs have functional suppressive effect on 3′ UTR reporters, despite that the small overlap between CLASH data and the pilot miRNA targeting map prevents us from deriving statistical estimate of the percentage of functional CLASH sites. Consistent with this notion, even though miRNA binding sites were frequently observed in coding sequences [6,7], it is believed that the majority of these interactions do not lead to strong target regulation [2,11]. This notion gained additional support from a recent study using PAR-CLIP on human embryonic stem cells, where only ∼21% of detected target gene RNA expression could be altered when the corresponding miRNA is perturbed [31], although some level of ambiguity of computational assignment of miRNA:target relationship with the PAR-CLIP method complicates the interpretation of these data.

Several lines of evidence support the quality of the data in our pilot miRNA-targeting map. First, the variation for the majority of the reporter assays is small. Second, known repressive miRNA:3′ UTR relationships have been captured. Third, both reporter assays in HeLa cells and western validation experiments in relevant cell types indicate that we can capture effects of miRNAs on genes not expressed in 293T cells. Fourth, known rules for miRNA target recognition were supported by our data. The Western blotting data also support that the false-positive rate of our data is low. We noticed that a reported suppressive effect of miR-145 on POU5F1 and SOX2 [32] cannot be observed in our pilot miRNA-targeting map. Neither were we able to observe the miR-145-based POU5F1 and SOX2 protein repression in NCCIT cells despite a strong overexpression of miR-145. Note that we did not use human embryonic stem cells as used in the original report [32]. This is because miR-145 can cause embryonic stem cell differentiation [32], and it will be difficult to distinguish a direct effect of miR-145 on POU5F1 and SOX2 versus an indirect effect via miR-145-induced embryonic stem cell differentiation. Although we do not fully understand why our data are different, we did notice that a CLIP study in human embryonic stem cells did not identify miR-145 binding on POU5F1 or SOX2 mRNA [31].

In the computational work, we utilized the high-throughput reporter data in the development of a new quantitative total score for modeling the total regulatory effects by both seed and seedless sites, as well as component scores for assessing individual contributions by seed sites and seedless sites. We adopted two assumptions in our modeling efforts: the linear additivity of contributions from multiple sites, and the hierarchical order in efficacy by different site types. During the model development process, we derived efficacy parameter estimates for modeling the variation in the effectiveness of different site types. To assess the predictive value of the total score, we analyzed high-throughput functional data from mRNA expression and proteomics studies. We found that, each of the scores can discriminate the potent miRNA inhibition from the weak inhibition and is thus useful for quantitative prediction of the levels of downregulation. In particular, the inclusion of the contribution of seedless sites is important for modeling and prediction.

## Methods

### Cell culture

293T cells and HeLa were cultured in DMEM (Invitrogen) with 10% FBS (Invitrogen) and 1% Penicillin-Streptomycin-Glutamine (Invitrogen). NCCIT and K562 cells were cultured in RPMI (Invitrogen) with 10% FBS and 1% Penicillin-Streptomycin-Glutamine. All cells were cultured at 37°C with 5% CO_2_.

MiRNA overexpression retrovirus was produced as previously described (Lu et al. 2008) by co-transfecting 293T cells with miRNA constructs and retroviral gag, pol, and VSVG protein expression constructs. Viral infection was carried out using spin-infection by centrifuging cells, retrovirus, and 8 µg/ml polybrene (Millipore) in a retronectin (Clontech)-coated well of a 6-well plate. Cells were selected with 5 µg/ml puromycin.

### Constructs

The miRNA expression library used in this study comprises 461 single miRNAs or miRNA clusters, each derived from human genomic DNA through cloning. MiRNA overexpression constructs and their cloning were previously described [23,27]. Briefly, genomic fragments containing miRNA hairpin and flanking sequences were cloned into a retroviral expression construct pMIRWAY-puro [25,27]. The control construct for miRNA is the vector backbone without any miRNA insert. These constructs were also used to produce virus to examine their effects on endogenous proteins.

3′ UTR dual-luciferase reporter constructs were cloned from human genomic DNA (except for Bmi1, which was cloned from mouse genomic DNA) into Promega’s psiCHECK2 vector through PCR. The specific enzyme sites, along with the start and end sequences of each 3′ UTR are listed below. For the 3′ UTR control, the unaltered Promega psiCHECK2 was utilized.

POU5F1 (XhoI & NotI) ggtgcctgcccttctaggaa…acagtagatagacacactta; SOX2 (XhoI & NotI) gggccggacagcgaactgga…ttgaaatatggacactgaaa; NANOG (XhoI & NotI) agatgagtgaaactgatatt…gtaaatatacagcttaaaca; KLF4 (XhoI & NotI) atcccagacagtggatatga…aacctataatattttatctg; LIN28 (XhoI & NotI) gccacaatgggtgggggcta…tattggtacgcaaactgtca; MYC (XhoI & NotI) ggaaaagtaaggaaaacgat…aaatatatcattgagccaaa; TP53 (PmeI & NotI) acattctccacttcttgttc…acctgtgtgtctgaggggtg; EIF2C2 (XhoI & NotI) catgttttagtgtttagcga…tatatatctggtttgcagta; DICER1 (XhoI & NotI) aaccgctttttaaaattcaa…ataaagttatcgtctgttca; MYB (XhoI & NotI) acatttccagaaaagcatta…gatgtgcccttattttacct; Bmi1 (PmeI & NotI) actgttaaggaaaagatttt…taataaaacaacagaaagat.

### High-throughput luciferase reporter assay

293T cells were trypsinized and resuspended at a concentration of 150,000 cells/ml in culture media. We used robotics in cell seeding and transfection, as detailed below, but such steps can also be carried out manually. The CyBi-Well automatic liquid handler (CyBio) was used to seed 20 µl of the cell dilution (3,000 cells) into each well of white, clear-bottom, 384-well plates (Corning 3707). The plates were then spun at 453 *g* for 30 seconds and cultured overnight. For experiments evaluating the density of cells, we varied the concentration of the cell suspension before seeding.

The following day, transfection mixtures were prepared in 96-well PCR plates. We included two types of controls on each plate, including control miRNA assayed with each of the 3′ UTR reporters (including a control reporter), and a control reporter assayed with each of the miRNAs on the plate (including a control miRNA construct). DNA mixture for each pair of 3′ UTR reporter and miRNA construct was prepared in each well to contain 27 ng of 3′ UTR reporter and 243 ng of miRNA construct in 5.4 µl total volume). For each well, 0.81 µl FuGENE 6 (Promega E2692) and 11.79 µl serum-free DMEM were mixed as per the manufacturer’s instructions. We prepare the DMEM/FuGENE6 master mixture in a microcentrifuge tube for all the wells on a 96-well DNA plate, and then use multichannel pipet to add 12.6 µl of the mixture into each DNA mixture well. The plate was spun at 453 *g* for 30 seconds and incubated at room temperature for 15 minutes. Using the automatic liquid handler, each transfection mixture was then added to the 293T cells seeded in the 384-well plates at 4 µl per well in quadruplicate, with each well receiving a transfection of 6 ng of 3′ UTR reporter, 54 ng of a miRNA construct, and 0.18 µl FuGENE 6 in 2.61 µl serum-free DMEM. Plates were then cultured for 48 hours or a specified amount of time.

Dual luciferase assays were performed using the Dual-Glo Luciferase Assay System kit (Promega E2940) as per the manufacturer’s instructions. To avoid signal variation between plates, we planned the experiments so that each plate experienced the same amount of time between adding luciferase reagent and plate reading. Luminescence levels were read out on a Wallac Victor^2^ plate reader (Perkin Elmer).

### Data preprocessing and normalization

Raw luciferase reporter data were preprocessed and normalized using custom Matlab codes. First, we calculated the ratio of *Renilla* luciferase activity (reflect 3′ UTR activity) versus firefly luciferase activity (control luciferase in the dual luciferase reporter), which is referred to as RvF ratios. Wells with weak readings were removed before the next steps of processing. Weak wells were defined as those with weak control firefly luciferase activity with a raw reading below 10,000. RvF ratios were then normalized based on the following formula. Normalized Luciferase Activity = (RvF_Well_ / mean(RvF_UTR-CtrlMiR_))/(mean(RvF_CtrlUTR-miR_)/mean(RvF_CtrlUTR-CtrlMiR_)). RvF_Well_ represents the RvF ratio of an assay, with mean (RvF) referring to the mean readings from all replicates of the corresponding control wells on the same plate as the RvF_Well_ assay. UTR-CtrlMiR stands for test 3′ UTR reporter assayed with a control miRNA construct. CtrlUTR-miR stands for a control 3′ UTR reporter assayed with a test miRNA. CtrlUTR-CtrlMiR stands for a control 3′ UTR reporter assayed with a control miRNA construct. After normalization, the mean of control assays becomes 1.

To identify miRNA-3′ UTR interactions that resulted in ≥25% downregulation, we examined the mean luciferase activity after normalization. Similarly, mean luciferase activities were applied to other cutoffs used in this study.

For each miRNA, we assigned major and minor strands based on miRbase data [33] and in-house next-generation sequencing reads. We analyzed predictions with both major and minor strands, as well as only with major strands.

To compare to CLASH dataset, CLASH data were downloaded from the supplement of Helwak *et al.* [8]. Only 3′ UTR binding sites were used to compare to miRNA-targeting map data.

### Western blot analysis

Western blot analyses were performed as described previously [27]. For NCCIT cells, collections were conducted on ice by scrapping in cold PBS. Cells were lysed using RIPA buffer supplemented with protease inhibitors (Roche 11873580001) per manufacturer’s instructions on ice for 20 minutes. Samples were then sonicated and spun at 16,100 *g* for 30 minutes. The supernatant was transferred to a new microcentrifuge tube, and protein was quantified by Bradford Assay (BioRad). Samples were boiled at 95°C in SDS loading buffer for five minutes, and were analyzed using 10% polyacrylamide gels. For K562 cells, cells counts were determined, and cells were then directly boiled in SDS loading buffer, and analyzed using 10% polyacrylamide gels. Samples were loaded based on equal cell counts. We obtained more quantitative data with this method in K562 cells. Antibodies used were Oct-4A (C30A3C1), Sox2 (D6D9), Nanog (D73G4), and Lin28A (A117) from Cell Signaling Technology, and Myb (clone 1-1) from Millipore.

To quantify the western data, we exclusively considered non-saturated exposures. In addition, quantification was performed on several non-saturated exposures with different exposure times and band intensities. To ensure data accuracy, each exposure was quantified using ImageJ software, and the relative protein amounts were calculated relative to loading control. Similar results were obtained with different exposures.

### RNA expression analysis

The miRNA expression profiles of 293T cells were determined using a plate-capture-based labeling and a bead-based detection system, as described before [34,35]. Data normalization was performed as described [35].

The expression levels of mature miRNAs in other experiments were determined by quantitative RT-PCR. Total RNA was collected using TRIzol Reagent (Invitrogen) as per manufacturer’s instructions. cDNA was prepared using the miScript II RT Kit (Qiagen), and qPCR was conducted with the miScript SYBR Green PCR Kit (Qiagen) as per manufacturer’s instructions. The Qiagen miRNA specific primers used during qPCR reaction were MS00003528 (miR-145), MS00004060 (miR-371), MS00003423 (miR-125a), and MS00006517 (miR-16). Relative gene expression was calculated using the ddCt method and human RNU6B for normalization.

### Statistical analysis

Student’s t-test was used to evaluate the significance of luciferase reporter data and expression data.

### STarMir prediction of miRNA binding sites

We analyzed the data from the 3′ UTR study. Rvf ratio, a measure of the level of miRNA-mediated 3′ UTR regulation relative to a baseline control, is computed from the readouts of the reporter assay. An Rvf ratio of around one indicates a lack of regulation. For ratios below one, a lower ratio indicates a greater level of down-regulation.

We first performed the prediction of miRNA binding sites using the STarMir program [30] for all the miRNA:3′ UTR pairs in the luciferase dataset. Similarly, for data from high throughput mRNA expression and proteomics data [29], we performed prediction of miRNA binding sites for 2406 targets down-regulated at the mRNA levels and 1,635 targets down-regulated at the protein level. This yielded a list of various miRNA site types, including 8mer, 7mer, 6mer (regular or offset) [2], and seedless sites for each miRNA:target pair.

### Quantitative modeling

#### Site-type specific efficacy parameters

Usually, an 8mer seed site is more effective than a 7mer site, which is more effective than a 6mer or a seedless site [2]. To quantify such functional variation solely due to various site types (independent of site/context features), we introduce parameter α_site-type_ as the average efficacy of a specific site-type for target down-regulation, where site-type can be 8mer, 7mer, 6mer, or seedless, and the following constraint is imposed: α_8mer_ > α_7mer_ > α_6mer_ > α_seedless_.

A range of logistic probability thresholds was considered from 0 to 1.0 with an increment of 0.05, with the hope that we can get estimates under the above constraint. From the luciferase data, we assembled 471 3′ UTR: miRNA pairs with RvF below 1.0, a threshold considered adequate for ensuring down-regulation. This down-regulation dataset can be used to estimate the efficacy parameters. Specifically, we divide the dataset into four subsets. Set 1 includes 3′ UTR: miRNA pairs for which only seedless miRNA binding sites are present (i.e., no 6mer, 7mer, or 8mer sites); set 2 is for 6mer containing 3′ UTR:miRNA pairs, with or without seedless sites (no 7mer, 8mer sites); set 3 is for 3′ UTR:miRNA pairs with 7mers, with or without 6mer or seedless sites (no 8 mer sites); set 4 is for 8mer sites possessing 3′ UTR:miRNA pairs, with or without other site types. Note that a lower RvF ratio shows a higher level of down-regulation, thus a measure of regulatory efficacy can be given by as (1-RvF). For n1_seedless_, the total seedless sites in set 1, and total efficacy 1 as the sum of (1-RvF) for all 3′UTR: miRNA pairs in set 1, α_seedless_, the average efficacy per seedless site, can be estimated from the equation, α_seedless_ * n1_seedless_ = total efficacy 1, i.e., α_seedless_ = total efficacy 1/ n1_seedless_. Similarly, α_6mer_ * n2_6mer_ + α_seedless_ * n2_seedless_ = total efficacy 2, where n2_6mer_ is the total number of 6mer sites, n2_seedless_ is the total number of seedless sites, and total efficacy 2 is the sum of efficacy for this set. With the estimate available for α_seedless_, an estimate for α_6mer_ can be computed from the linear equation, i.e., α_6mer_ = (total efficacy 2 – α_seedless_ * n2_seedless_)/ n2_6mer_. Next, α_7mer_ * n3_7mer_ + α_6mer_ * n3_6mer_ + α_seedless_ * n3_seedless_ = total efficacy 3, where for set 3, n3_7mer_ is the total number of 7mer sites, n3_6mer_ is the total number of 6mer sites, n3_seedless_ is the total number of seedless sites, and total efficacy 3 is the sum of efficacy for this set. With estimates available for α_seedless_ and α_6mer_, an estimate for α_7mer_ is calculated by (total efficacy 3 – α_6mer_ * n3_6mer_ – α_seedless_ * n3_seedless_)/ n3_7mer_. Finally, α_8mer_ * n4_8mer_ + α_7mer_ * n4_7mer_ + α_6mer_ * n4_6mer_ + α_seedless_ * n4_seedless_ = total efficacy 4, where for set 4, n4_8mer_, n4_7mer_ and n4_6mer_ are total numbers of 8mer, 7mer and 6-mer sites, respectively, and total efficacy 4 is the sum of efficacy for this set. With estimates available for α_seedless_, α_6mer_ and α_7mer_, an estimate for α_8mer_ is given by (total efficacy 4 – α_7mer_ * n4_7mer_ –α_6mer_ * n4_6mer_ – α_seedless_ * n4_seedless_)/ n4_8mer_.

#### A quantitative score

We aim to develop a score for quantitative prediction of the level of target down-regulation by miRNAs. The average efficacy parameters address functional variation among different site types. On the other hand, for sites of the same type, e.g., two 8mer seed sites, there can be significant differences in effectiveness due to other site/context features such as structural accessibility. This variation can be addressed by the logistic probability of individual site [22]. With the estimates of the efficacy parameters and logistic probabilities, for a target with *n* predicted sites of various types, we propose to compute a predictive score as a weighted sum of site probabilities: ∑*_site i_ Pr_i_* α_site-type_(*i*), where *i*= 1,…, *n,* and *Pr_i_* and α_site-type_(*i*) are the logistic probability and the site-type specific efficacy for the *i*th site on the target, respectively. This score not only addresses cooperative regulatory effects of seed and seedless sites, but also takes into account functional differences among different site types as well as differences due to other site features.

### Assessment of quantitative score

To assess the predictive value of the score, we analyze high throughput data from proteomics and microarray studies [29]. We compared the scores of the most down-regulated targets and the least down-regulated ones. Statistical significances were computed for the comparison.

## Acknowledgments

This work was supported in part by NIH grants 1R01GM099811, 1R01GM138856 (to Y.D. and J.L.) and 1R01CA149109 (to J.L.), NSF grant DBI-0650991 (to Y.D.), as well as a Connecticut Stem Cell Award to J.L. Additionally, S.M. was partially funded by the NIGMS/NIH-sponsored Predoctoral Training Grant in Genetics T32 GM007499. We thank Dianqing Wu and his lab for technical help.

